# Widespread loss of Y expression in the absence of transcriptional dosage compensation in *Rumex hastatulus*

**DOI:** 10.1101/2025.11.20.689509

**Authors:** Bianca M. Sacchi, Meng Yuan, Cassandre Pyne, Baharul Choudhury, Yunchen Gong, Spencer C.H. Barrett, Stephen I. Wright

**Affiliations:** Department of Ecology and Evolutionary Biology, University of Toronto, Toronto, Canada; Department of Biology, Queen’s University, Kingston, Canada; Centre for Analysis of Genome Evolution and Function, University of Toronto, Toronto, Canada

## Abstract

There is growing interest in the potential role of gene regulatory evolution in the expansion and degeneration of Y chromosomes. Evolutionarily young sex chromosomes, such as those found in the flowering plant *Rumex hastatulus*, can provide insights into the early stages of this process. Using a new high-quality genome assembly with a well-assembled Y chromosome and a highly replicated transcriptome dataset, we found widespread underexpression of Y genes compared to their homologous counterparts on the X chromosome. We also found evidence for increased expression differences for more distantly diverged gametologs, suggestive of progressive loss of expression of Y linked genes over time. Genes with greater expression loss on the Y chromosome also showed elevated rates of protein evolution, as expected if silencing alleles mask the effects of deleterious mutations on the Y and/or if selective interference drives early degeneration in regulatory sequences. However, in contrast with the predictions of recent models of sex chromosome expansion and degeneration by regulatory evolution, there was no evidence of early dosage compensation. Overall, we conclude that in *Rumex* Hill-Robertson interference alone may be the main driver of Y chromosome degeneration, although further understanding of the temporal order of regulatory changes could help further untangle cause and effect.

## Introduction

Degeneration of the Y (and W) chromosome has been observed across many taxa and independent events in sex chromosome evolution (Abbott et al. 2017; Bachtrog 2013; Bergero et al. 2015; Bachtrog et al. 2008; Moraga et al. 2025). Degeneration describes processes such as deleterious mutation accumulation, gene loss, transposable element accumulation, and in the most extreme case, loss of the Y chromosome (Bachtrog 2013). The Y chromosomes of some taxa such as those in mammals and *Drosophila melanogaster* are highly degenerated and lacking in most ancestral protein-coding genes. Long-standing theoretical work predicts several causes and outcomes of sex chromosome evolution (Abbott et al. 2017). Sex chromosomes arise when recombining autosomes acquire a sex-determining factor. Theory predicts that linkage of sexually antagonistic variants to a sex-determining locus results in a selective advantage for strong linkage between these sites, driving further expansion of the sex-linked region. Deleterious mutation accumulation then follows driven by several potential mechanisms associated with selective interference among sites in a non-recombining region, including Muller’s ratchet, background selection and selective sweeps (Orr and Kim 1998; Bachtrog 2013; Charlesworth 1978; Hill and Robertson 1966; Rice 1987). In general, models of selective interference describe degeneration as a process that results from inefficient selection once recombination stops, leading to a decline in male fitness during Y degeneration.

Recent theoretical work has introduced an alternative mechanism for Y chromosome expansion and subsequent degeneration that does not necessarily require pre-existing sexual antagonism or selective interference (Lenormand and Roze 2022). First, the spread of recombination suppression on the Y chromosome is favoured due to drift-mediated regulatory divergence between X and Y gametologs, coupled with the rapid evolution of sex-specific dosage compensation (DC). Cis-regulatory loss of expression on the Y chromosome under this model acts as a dominance modifier, enabling deleterious mutation accumulation in a gene-by-gene fashion due to the masking effects of silencing alleles. The Lenormand and Roze (2022) model predicts a positive feedback loop where decreased Y expression allows deleterious mutations to accumulate in Y-linked genes, driving further loss of Y expression and ultimately silencing, even in the absence of selective interference. Thus, regulatory coevolution on X- and Y-linked genes could potentially accelerate Y degeneration and drive the expansion of sex-linked regions.

Several key predictions arise from models of sex chromosome evolution by regulatory evolution. First, asymmetric loss of expression of Y-linked gametologs should arise early, following sex linkage. Second, due to the masking effects on fitness deleterious mutations should accumulate more readily on Y gametologs that have reduced expression. Similarly, models of adaptive silencing would predict selection for reduced Y gametolog expression in genes with greater deleterious mutation accumulation (Crowson et al. 2017). Finally, the models predict the evolution of dosage compensation during early stages of sex chromosome expansion and degeneration, rather than DC evolving at a later stage in the process, provided stabilizing selection for total gene expression is sufficiently strong (Lenormand et al. 2020). Early evolution of dosage compensation is also key for regulatory divergence to drive the spread of recombination modifiers and the expansion of the sex-linked region (Lenormand and Roze 2022). Examination of the early stages of sex chromosome expansion and degeneration should provide important opportunities to test the predictions of this model, by investigating which changes occur early vs. later in the process of sex chromosome evolution.

Evolutionarily young sex chromosomes are valuable candidates for investigating the earliest processes and changes that occur during sex chromosome evolution (Charlesworth 2019). Very old sex chromosomes, such as those of placental mammals which arose at least 166 million years ago (Veyrunes et al. 2008), or the >60MYA ancestral sex chromosomes of *Drosophila*, are already highly degenerated (Bachtrog 2013). Although young neo-sex chromosomes such as those found in *Drosophila* provide important insights into the dynamics of early sex chromosome evolution, there is evidence that rapid chromosome-wide dosage compensation on these young sex chromosomes is driven in part by the recruitment of ancestral DC mechanisms such as the male-specific lethal complex (Zhou et al. 2013; Ellison and Bachtrog 2019).

Plants provide excellent model systems to investigate the processes that occur following the earliest evolution of sex chromosomes after the evolution of dioecy from hermaphroditism. Sex chromosome ages in plants tend to be evolutionarily young, diverging from <5 MYA to nearly 30 MYA (Prentout et al. 2021; Harkess et al. 2015, 2017; Charlesworth 2019; Janousek et al. 2022; Krasovec et al. 2018; Renner and Müller 2021). By investigating patterns of gene loss and expression divergence in such young sex chromosomes it should be possible to get a clearer picture of the timing of these events, which in turn can lead to a better understanding of the processes driving early sex chromosome evolution.

Studies to date in plants using transcriptome-based approaches have provided evidence for a directional loss of gene expression on Y-linked gametologs in species with large heteromorphic sex chromosomes, with mixed evidence for dosage compensation (Muyle et al. 2022). Early transcriptome studies demonstrated Y-linked expression loss in plants, however without genomic data, it can be difficult to distinguish gene silencing from loss (Bergero et al. 2015). Genomic data allows for a better estimation of gene loss vs. expression loss; for example in *Silene latifolia* it was estimated that approximately 45% of Y-linked genes in the sex-linked region are not expressed (Papadopulos et al. 2015). More recent work in this system shows decreased Y expression relative to X-linked genes overall (Moraga et al. 2025), and evidence for dosage compensation in many silenced genes, but only in half of the hemizygous genes (Chibalina and Filatov 2011; Muyle et al. 2012; Papadopulos et al. 2015). In *Carica papaya*, Y gametologs show decreased expression particularly in the older 7 MYA strata (Liu et al. 2021), with preliminary evidence for partial dosage compensation in a handful of shared genes and hemizygous genes (Liu et al. 2021; Chae et al. 2021). Evidence for loss of expression in the ∼5-10 million-year-old Y chromosome has also been observed in *Rumex hastatulus* using the allelic coverage of sex-linked SNPs in RNAseq data (Hough et al. 2014). However, in this study there was no evidence for dosage compensation on hemizygous and silenced genes. In *R. rothschildianus,* which has a different and potentially older sex chromosome origin than *R. hastatulus*, Y alleles were overexpressed in some male-biased genes; however, there was an overall trend of decreased Y/X allelic expression in up to half of identified gametologs. A subset of the putatively hemizygous and silenced genes in *R. rothschildianus* also exhibited signs of dosage compensation (Crowson et al. 2017). Therefore this growing body of evidence suggests that Y gametolog expression loss may occur early in sex chromosome evolution but may or may not be associated with a widespread dosage compensation mechanism.

Until relatively recently, studies investigating the expression of X-Y gametologs relied on transcriptome and/or genome assemblies from females owing to difficulties in sequencing Y chromosomes, which are often repeat-rich and can have low divergence with the X chromosomes. One potential challenge with inferring expression divergence by aligning male samples to a female reference is mapping bias; for example, if short transcriptome reads from the X chromosome map to the X chromosome reference at a higher rate than Y chromosome reads, this will lead to inaccurate estimates of the relative expression of Y-linked genes (Degner et al. 2009), the degree to which should correlate with X-Y divergence. Furthermore, transcriptome-based approaches make it difficult to distinguish gene loss from silencing of genes on the Y chromosome. Indeed, preliminary analyses of genomic coverage in *R. hastatulus* suggested that gene silencing may often precede loss (Beaudry et al. 2017); but our understanding of the temporal dynamics of gene silencing and loss remain incomplete in the absence of high-quality Y chromosome assemblies. Advances in long-read sequencing and chromosome conformation capture allow for improved assemblies of X and Y chromosomes in their entirety (Carey et al. 2021; Moraga et al. 2025; Sacchi & Humphries et al. 2024). Analysis of fully phased sex chromosomes can provide a more complete picture of early regulatory evolution and gene loss on the sex chromosomes resulting in a better understanding of the timing of regulatory divergence, gene loss and deleterious mutation accumulation.

Here, we examine expression evolution on the sex chromosomes of the wind-pollinated herb *R. hastatulus* (Polygonaceae) to address several key questions regarding the role of regulatory divergence in sex chromosome formation and degeneration. Generating a new fully phased genome assembly of a male from the XY (Texas) cytotype of this species, we use a large, highly replicated population sample (75 males, 74 females) from this cytotype to quantify expression divergence between the sexes and the sex chromosomes. This high level of replication across multiple genotypes enabled an unusual level of statistical power to explicitly test for dosage compensation and changes in the expression level of Y and X gametologs, using genomic sequencing to control for mis-mapping of reads. We generated a new reference assembly and focused on the XY cytotype to reduce the complexities of mis-mapping of reads between the sex chromosomes in the extremely young neo-sex chromosomes of the XYY cytotype (assembly by Sacchi & Humphries et al. 2024). Using these data we address the following questions: 1) How rapidly does gametolog-specific expression evolve, and is it biased towards expression loss on the Y? 2) Is there evidence for early and rapid evolution of dosage compensation? and 3) Is there an association between Y expression loss and deleterious mutation accumulation? Addressing these questions should provide important evidence for evaluating the predictions of recent models of sex chromosome evolution by regulatory evolution.

## Methods

### Genome assembly details

We performed PacBio and Dovetail Omni-C sequencing (Lieberman-Aiden et al. 2009) on an *R. hastatulus* XY male following the protocol of Sacchi & Humphries et al. (2024). The sequenced male sample Rh15 was a result of an F_1_ cross between a male and female from independent maternal families collected from Rosebud, TX.

We quantified DNA samples using Qubit 2.0 Fluorometer (Life Technologies, Carlsbad, CA, USA). The PacBio SMRTbell library (∼20kb) for PacBio Sequel was constructed by Canatata Genomics using SMRTbell Express Template Prep Kit 2.0 (PacBio, Menlo Park, CA, USA) using the manufacturer’s recommended protocol. The library was bound to polymerase using the Sequel II Binding Kit 2.0 (PacBio) and loaded onto the PacBio Sequel II. Sequencing was performed on PacBio Sequel II 8M SMRT cells. Additional sequencing was performed by Cantata Genomics using PacBio Revio sequencing. DNA samples were quantified using Qubit 2.0 Fluorometer (Life Technologies, Carlsbad, CA, USA).

For each Dovetail Omni-C library, chromatin was fixed in place with formaldehyde in the nucleus. We digested fixed chromatin with DNase I and then extracted, chromatin ends were repaired and ligated to a biotinylated bridge adapter followed by proximity ligation of adapter containing ends. After proximity ligation, crosslinks were reversed and the DNA purified. Purified DNA was treated to remove biotin that was not internal to ligated fragments. We generated sequencing libraries using NEBNext Ultra enzymes and Illumina-compatible adapters and biotin-containing fragments were isolated using streptavidin beads before PCR enrichment of each library. The library was sequenced on an Illumina HiSeqX platform to produce ∼ 30x sequence coverage.

For the Revio sequencing, we quantified DNA samples using Qubit 4.0 (ThermoFisher, Waltham, MA), and these were assessed on Femto Pulse (Agilent, Santa Clara, CA) prior to library preparation. DNA samples that passed quality control were sheared on MegaRuptor 3 (Diagenode, Denville, NJ) to 15-18kb. Libraries were constructed using SMRTbell Prep Kit 3.0 (Pacific Biosciences, Menlo Park, CA) according to manufacturer protocol. Adaptor-ligated libraries were size-selected with AMpure PB beads to remove fragments less than 5kb, or on BluePippin instrument (Sage Science, Beverly, MA) to remove fragments less than 6 to 15kb, depending on available library material. Libraries were bound to polymerase with Revio Plolymerase Kit, then sequenced with Revio SMRT cell (25M) on the Revio instrument (Pacific Biosciences, Menlo Park, CA) for 24 hours.

We generated a haplotype-resolved de novo assembly using Hifiasm v. 0.16.1-r375 (Cheng et al. 2021) and the Omni-C sequence for phased haplotype resolution. Paired-end Omni-C reads were mapped, then filtered to the two phased assemblies using bwa v0.7.15 (Li and Durbin 2009) following the Arima mapping pipeline (https://github.com/ArimaGenomics/mapping_pipeline), and the resulting filtered (MapQ > 10) bam files had duplicates marked using Picard v2.7.1. We scaffolded both haplotypes of the assembly using YAHS v1.2a.2 (Zhou et al. 2023) to generate scaffolded assemblies from each phased haplotype. To construct contact maps we used BWA v0.7.15 to align Omni-C reads to our final assembly. We then used pairtools v1.0.2 to create a pairsam file with annotations of ligation events and potential pairs (Open2C et al. 2023). The minimum threshold used for defining multimapping alignments was 40 and the maximum gap between alignments was 30.

We then sorted the parsed pairsam file and marked duplicate pairs using pairtools (pairtools sorted and dedup). The pairs were then split into a bam and pairs file using pairtools split. Lastly, we converted the pairs to contact maps using Juicer’s ‘pre’ command (Durand et al. 2016b). All contact maps were visualized using the Juicebox visualization environment (Durand et al. 2016a).

We manually inspected the scaffolded assembly using a combination of Juicebox v1.11.08 (Durand et al. 2016a) and comparisons between the two haplotypes and with our previous assemblies, supported by genetic maps (Rifkin et al. 2022, 2021). Based on manual inspection and using the AGP files, we identified and corrected several false joins: in the maternal haplotype, scaffold 1 was split into the X chromosome and A2, and scaffold 3 into A3 and A4. In the paternal haplotype, scaffold 3 was split into A3 and A4. Assembly completeness was assessed using BUSCO v5.4.4 (Manni et al. 2021) using the Eukaryota databases. BUSCO scores and scaffold N50, L50, N90 and L90 statistics are available in Supplementary Table 1.

### Gene annotation

Gene annotation followed previous approaches (see Rifkin et al. 2022). We performed the annotation with MAKER-3.01.03 (Cantarel et al. 2008) in four rounds. In the first round, we used the *R. hastatulus* RNA-Seq transcripts from previously published floral and leaf transcriptomes (Hough et al. 2014; Sandler et al. 2018) and annotated Tartary buckwheat proteins from version FtChromosomeV2.IGDBv2 (Zhang et al. 2017) for inferring gene predictions, and used a transposable element (TE) library generated by RepeatModeler (Smit and Hubley 2008) to mask the genome. We trained the resulting annotation for SNAP gene predictor, using the gene models with an AED of 0.5 or better and a length of 50 or more amino acids. In the following rounds, we used the resulting EST and protein alignments from the first round, and the SNAP model from the previous round for annotation. We functionally annotated the final gene models based on BLAST v 2.2.28+ (Altschul et al. 1990) and InterProScan 5.52–86.0 (Jones et al. 2014), by using the related scripts in the Maker package.

### Sex-linked SNP identification

We mapped RNAseq leaf expression data from 12 population samples of XY cytotype male and female plants of *R. hastatulus* (Hough et al. 2014) to both haplotype assemblies using STAR v2.7.10a (Dobin et al. 2013). We performed variant calling using freebayes v1.3.4 (Garrison and Marth 2012) and then filtered sites to a final set comprised of biallelic sites with genotype quality > 20. We then used custom scripts to identify putative sex-linked SNPs. We selected all sites that were heterozygous in all six males and homozygous in all six females to obtain candidate fixed SNP differences between X and Y. Using this approach, we confirmed the sex chromosomes and inferred that the pseudoautosomal region occupies the first approximately 45 Mb of the Y chromosome and the terminus of the X chromosome assembly starting at approximately 220 Mb (Supplementary Figure 2).

### Population description and sequencing details

*R. hastatulus* RNA and DNA samples from field-collected maternal families were collected and processed as previously described (Yuan et al. 2025). In summary, *R. hastatulus* samples were collected from a population in Rosebud, Texas, US (Pickup and Barrett 2013) and seedlings planted in a glasshouse at the Earth Sciences Center, University of Toronto. These population samples underwent a generation of random mating from independent maternal families, and the F_1_ generation was then grown in the glasshouse. Leaf tissue was collected from male and female F1 plants for DNA and RNA isolation. The standard protocols of Qiagen Plant Mini kit and Sigma Aldrich Spectrum Plant Total RNA Kits for DNA and RNA isolation were used, respectively. All libraries were prepared and sequenced at the Genome Quebec Innovation Centre in Montréal, Canada. The DNA samples were sequenced at a depth of 10-15X. 75 males and 74 female samples were used for this study.

### Read mapping

To facilitate read mapping, we created a reference genome from the haplotype assemblies that contains the X chromosome (with no pseudoautosomal region, or PAR), the Y, the Y pseudoautosomal region, and one copy of each autosome (from the paternal haplotype assembly). This assembly allowed us to use a competitive mapping approach to determine read counts for X and Y gametologs within the sex-linked regions, while allowing for unique mapping of PAR and autosomal regions. Genomic reads for 75 male and 74 female XY *R. hastatulus* leaf samples were mapped to the reference described above using bwa-mem2 v2.0pre2, and duplicates were marked using samtools v1.20. RNA reads for 75 male and 74 female samples were mapped to the same reference using STAR v2.7.10a in two-pass mode with default parameters. Female samples were mapped to a reference containing the X and Y chromosomes to better identify mismapping of female reads to the Y chromosome, which likely results from high sequence similarity between the sex chromosomes.

### Identification of X-Y gametologs and hemizygous genes

One-to-one gametologs shared between the X and Y chromosomes were identified using GENESPACEv1.3.1 (Lovell et al. 2022), which uses dependencies Orthofinder v2.5.5 (Emms and Kelly 2019) and MCScanX (Wang et al. 2012). Genespace was run on the two complete haplotype assemblies of the XY cytotype, along with the hermaphroditic outgroup *R. salicifolius* (Sacchi & Humphries et al. 2024). The list of gametologs includes both syntenic and non-syntenic orthologs, and we included both, as long as they had a one-to-one gene copy relationship between X and Y. X-hemizygous genes were identified as follows: if an X-linked gametolog is present in the outgroup *R. salicifolius*, but absent from the Y in the GENESPACE output, it was considered putatively hemizygous. This is a conservative estimate of the number of genes lost, as *R. salicifolius* is the most distantly related hermaphroditic species sequenced in the clade (Hibbins et al. 2024). Following the method from Sacchi & Humphries et al. 2024 for the XYY cytotype, we performed a BLAST nucleotide search (v2.2.31+) of putatively hemizygous X chromosome genes to the haplotype-level genome assembly containing the Y chromosome. Genes with greater than 50% alignment to the Y chromosome were removed from the list of hemizygous genes. Genes with <50% alignments were considered putatively partially lost genes, whereas genes with no blast hits to the Y chromosome were categorized as completely lost from the Y.

### RNA and DNA read counting

We performed read counting on both DNA (as a control) and RNA mapped to the combined reference genome assembly bearing X, Y, and one copy of each autosome for the *R. hastatulus* XY cytotype using featureCounts v2.0.2 parameters “-t exon -g gene_id” and excluding multiply-mapping reads. This reference genome included a single copy of the pseudoautosomal region (PAR), to avoid cross-mapping of PAR reads to both X and Y. Due to high sequence similarity between some X-Y gametologs, filtering was necessary to remove gametologs where reads from the X chromosome of female samples mapped excessively to the Y chromosome, as these genes will also likely have skewed estimates of X/Y expression levels in males. First, Y genes with a mean read count >20 across all female samples in either the RNAseq and/or genomic data were removed. The associated X gametologs for these mismapping Y genes were also removed from further analysis.

Additionally, we identified genes with mapping bias by estimating gametolog-specific differences in genomic read counts in males. We modified the DESeq2 (Love 2017; Love et al. 2014) analysis pipeline to calculate gametolog-specific read counts, transforming the original read count data frame so that each row is a X-Y gametolog, and each column contains the X and Y counts for each male sample, separately. Genes with significant differences between X and Y gametolog read counts in the male genomic DNA were removed before gametolog-specific expression RNAseq analysis (FDR adjusted p-value <0.1, |log2FC| > 0.5), as this suggested mapping bias of genomic DNA, where an equal read ratio between X and Y is expected. Together, these two approaches allowed us to eliminate genes with significant mapping biases affecting our estimates of expression divergence between the sex chromosomes. This, however, does lead to a significant drop in the number of X-Y genes we can analyze for gametolog-specific expression (Supplementary Table 2).

For our dosage compensation analysis, we ran DEseq2 on each sample (male and female) with and without sex-linked genes included. We used the output from the analysis without sex-linked genes to obtain normalization factors based on library size and used these factors to normalize the read counts of the entire dataset, ensuring that normalization did not obscure real differences in expression due to different amounts of sex-linked reads between male and female samples. Comparison of the DESeq size factors with and without the sex-linked genes showed negligible differences (Supplementary Figure 4.)

### Gametolog-specific expression

To assess which gametologs have significant differences in expression levels between X and Y copies, we used an adapted DESeq2 approach (Love 2017; Love et al. 2014). Read counts did not need to be normalized, and all size factors were set to 1 since counts from each sample were separated into X and Y counts and sample ID was a factor in the design (∼sample + allele). We applied the Benjamini-Hochberg FDR correction to *p*-values for each gametolog and used a significance threshold of adjusted *p* < 0.1 to determine which gametologs showed differences in X vs. Y expression within the samples.

### Estimating synonymous and nonsynonymous substitution rates

To calculate branch-specific nonsynonymous (*dN*) and synonymous (*dS*) substitution rates, we fit the Standard MG94 model from the Hyphy standalone analyses suite (Muse and Gaut 1994), obtained from https://github.com/veg/hyphy-analyses, using HYPHY version 2.5.62(MP) (Kosakovsky Pond et al. 2020). To run this model, we first identified 1-1-1-1 orthologs shared between *R. bucephalophorus* (Hibbins et al. 2024) as the closest hermaphroditic outgroup, *R. sagittatus* (Hibbins, Pyne et al., in prep) as a more distant hermaphroditic relative and both *R. hastatulus* haplotype assemblies using GENESPACEv1.3.1 (Lovell et al. 2022). Then we used MUSCLEv3.8.1551 (Edgar 2004) and the hyphy standalone tools ‘pre-msa’ and ‘post-msa’ to perform codon alignment, and IQ-TREEv2.2.6 (Minh et al. 2020) to create unrooted maximum likelihood gene trees. We calculated the difference in *d_N_* between X and Y gametologs to estimate the difference in non-synonymous substitutions between the sex chromosomes. To estimate *d_S_* for the entire XY branch, as an estimate of time since the gametologs coalesced, we summed the X and Y *d_S_* estimates and filtered out genes with XY *d_S_* above 0.5 as these may not reflect true gametologs.

## Results

### Significant gametolog-biased expression shows widespread silencing of Y genes

After filtering for cross-mapping between X and Y gametologs, we analyzed 267 genes with shared one-to-one orthology between the X and Y chromosomes of *R. hastatulus* (Table B2). Of those gametologs, 111 (42%) show a significant difference (FDR adjusted *p*-value < 0.1) in gametolog read count across 75 male samples using a log2FC cutoff of 1 (Figure 1). Among these differentially expressed gametologs, there is a strong enrichment of genes with significantly reduced Y/X expression and this excess increases with an increasing fold-change cutoff (Table 1). Thus, we see an overall pattern suggesting Y expression loss.

**Figure 1.**
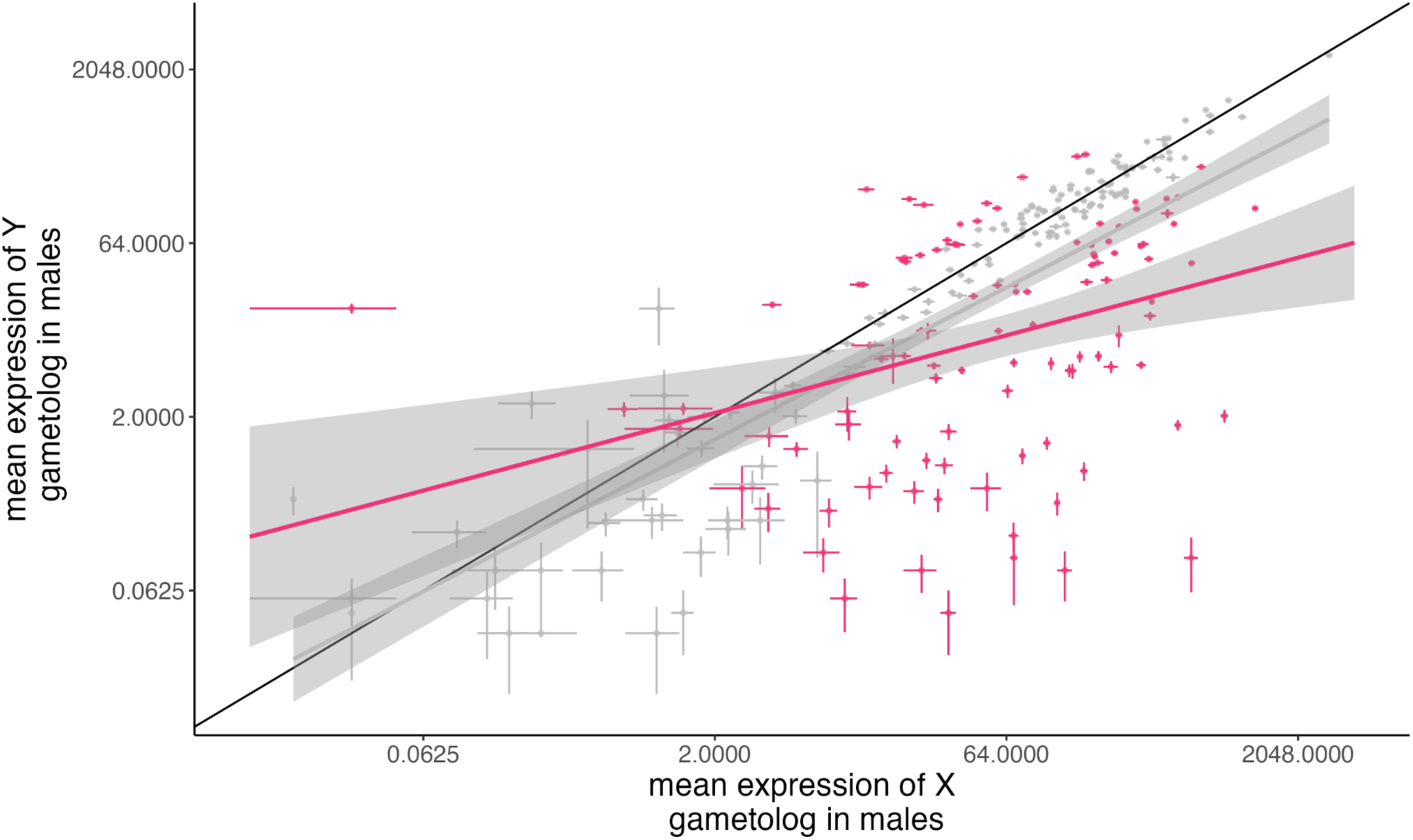
Mean expression of X-linked gametologs (*n*=267) compared to mean expression of the corresponding Y-linked gametolog in 75 *R. hastatulus* male plants. Points represent either gametologs with a significant difference in gene expression (FDR <0.1) and absolute log2 fold change >1 (magenta) (adj *R*^2^ = 0.1019, *p* = 0.00047). Error bars represent log2 fold change standard error across samples. The regression line in grey is for all the data points (significant and nonsignificant gametolog-specific expression) (adj *R*^2^ = 0.52, *p* <2.2e −16).

**Table 1.** Number of genes with significant (padj < 0.1) gametolog-specific expression at different log2 fold change cutoffs.

| Number of genes increased Y/X expression | Number of genes with decreased Y/X expression | Log2 fold change cutoff (absolute value) |
| --- | --- | --- |
| 47 | 103 | 0.5 |
| 27 | 84 | 1.0 |
| 13 | 70 | 1.5 |
| 9 | 59 | 2.0 |

### Divergence patterns show evidence for increased Y expression loss over time and possibly over strata

We estimated *d_S_* of the X-Y branch for each X-Y gametolog, as a proxy for the timing of sex linkage for each gene. There was a significant, albeit weak, negative association between Y/X read ratio (log2) and *d_S_* (Figure 2). As expected, clusters of low *d_S_* genes were present in the Y pseudoautosomal region which occupied 0-45Mb (excluded from the gametolog-specific expression analysis). Low *d_S_* clusters were also present around 100Mb and at the end of the chromosome (∼500Mb) (Figure 3) and may represent younger sex-linked strata. Genes with significant Y/X silencing tended to lie in less gene-dense regions and have elevated *d_S_* estimates compared to the background level. This finding is consistent with the prediction of greater Y expression loss in older, more degenerated sex-linked regions.

**Figure 2.**
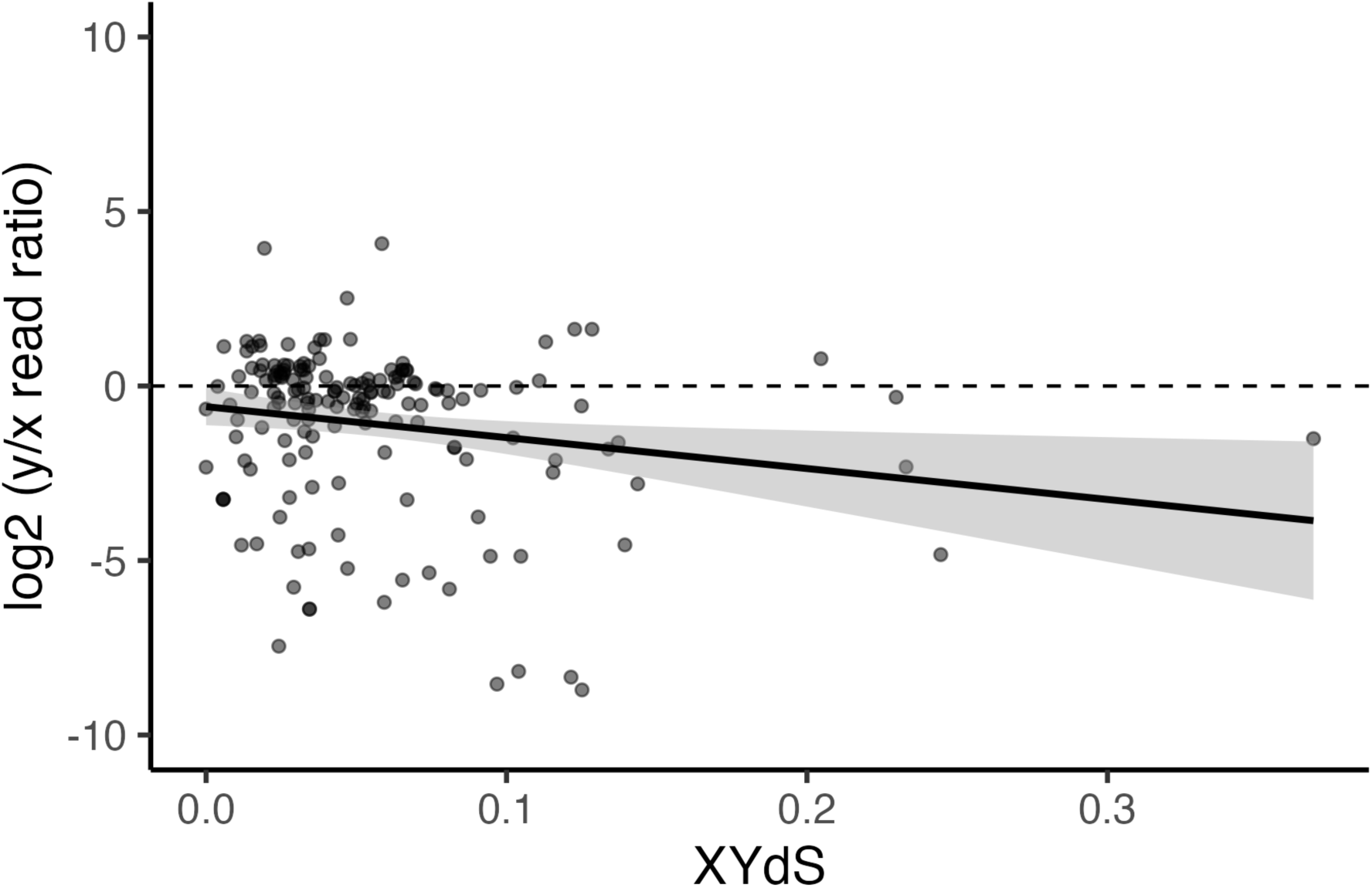
Linea relationship between log2(Y/X read ratio) and XYdS. Each point represents a gametolog pair. Read ratio is averaged across 75 male plants of *Rumex hastatulus* for each set of gametologs. The dashed horizontal line indicates where X and Y gametologs have equal expression. Adjusted *R*^2^ = 0.0294, *F*-statistic: 5.968 (1, 163)*, p*-value: 0.01564.

**Figure 3.**
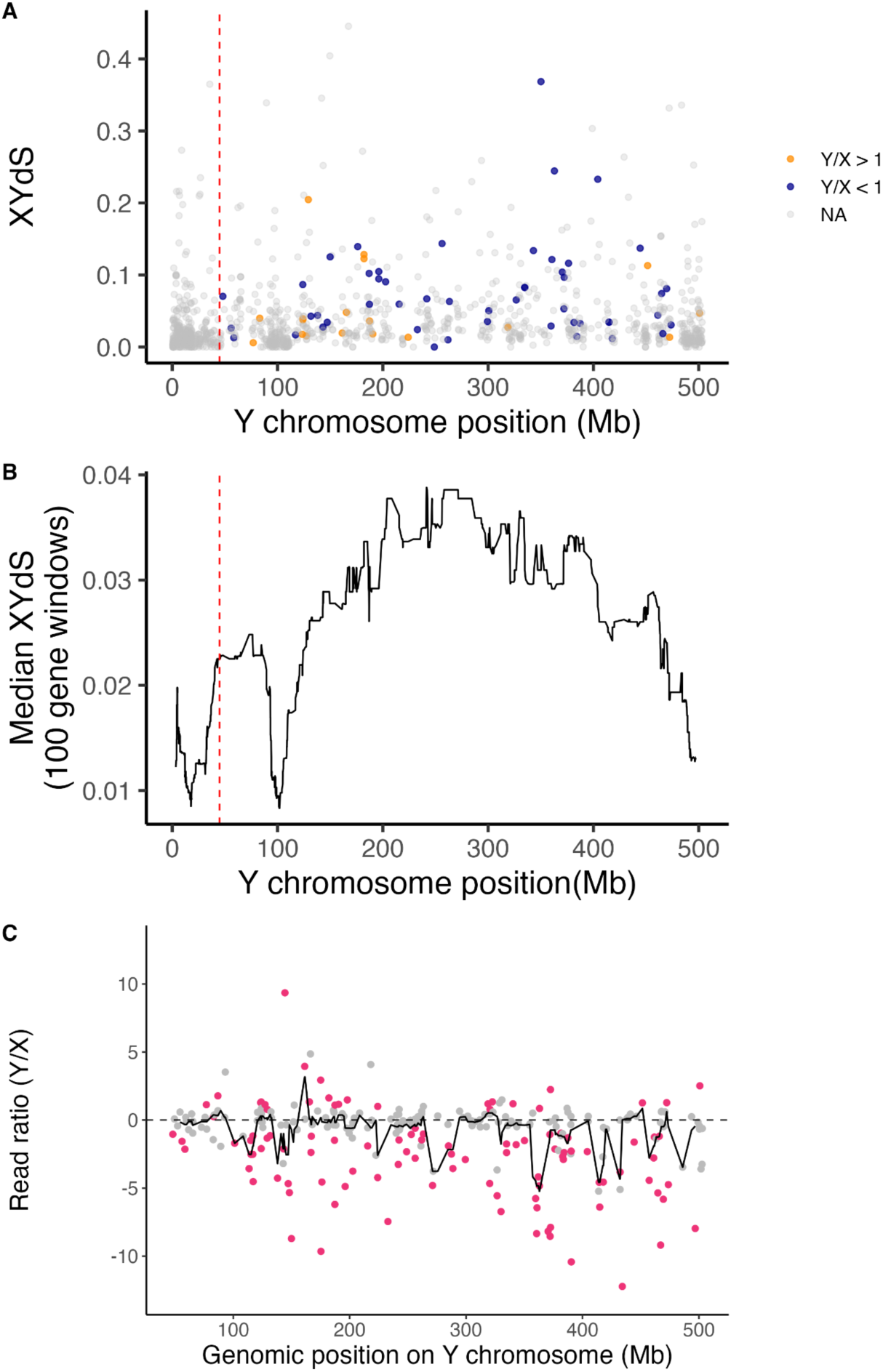
XY*dS* estimates relative to the genomic position on the Y chromosome of *Rumex hastatulus*. The dashed red line represents the end of the pseudoautosomal region and the beginning of the sex-linked region. A) XY*dS* estimates of X-Y gametologs, orange points represent gametologs where Y expression is significantly different and at least 2-fold greater than X expression, while blue points represent gametologs where Y expression is 2-fold lower than X expression. B) Median XY*dS* calculated in sliding windows of 100 genes with a step size of 1. C) b) Log2 of Y/X read ratio vs. Y chromosome position for each gametolog pair. Points below zero have decreased Y expression relative to X. Genes in magenta show significant gametolog expression (abs log2FC >1, *p* <0.1). The black line indicates the median log2 Y/X read ratio, computed in 10Mb sliding windows with a step size of 1Mb.

### Silenced Y genes show evidence of elevated molecular evolutionary rates

To test for greater deleterious mutation accumulation in genes with more Y expression loss (Lenormand and Roze 2022), we estimated branch-specific *d_S_* and *d_N_* for both the X and Y gametolog branches using HYPHY MG94Fit. We observed a significant negative relation between the log2-transformed Y/X expression ratio and difference in *d_N_* (‘*d_N_* difference’) between Y and X gametologs (Figure 4), consistent with the prediction that Y-linked genes with greater expression loss experience accelerated molecular evolution and higher genetic load. However, this may be due to an effect of XY*d_S_* on both *d_N_* difference and expression loss, due to the role of sex chromosome divergence time on both deleterious mutation accumulation and Y expression loss. To test if there was an effect of Y expression loss on deleterious load while controlling for *dS* we conducted the following model comparison: *d_N_* difference ∼ XY*d_S_* and *d_N_* difference ∼ XY*d_S_* + log2(Y/X ratio). The second model had the lowest AIC, suggesting that both XYdS and Y/X expression ratios predict dN difference (Supplementary Figure 7). Overall, these results suggest that lower Y/X expression ratios are correlated with a higher non-synonymous substitution rate in the Y gametolog compared to the X gametolog, controlling for time since sex linkage using *dS*. This result is consistent with our prediction that Y expression loss may contribute to deleterious mutation accumulation.

**Figure 4.**
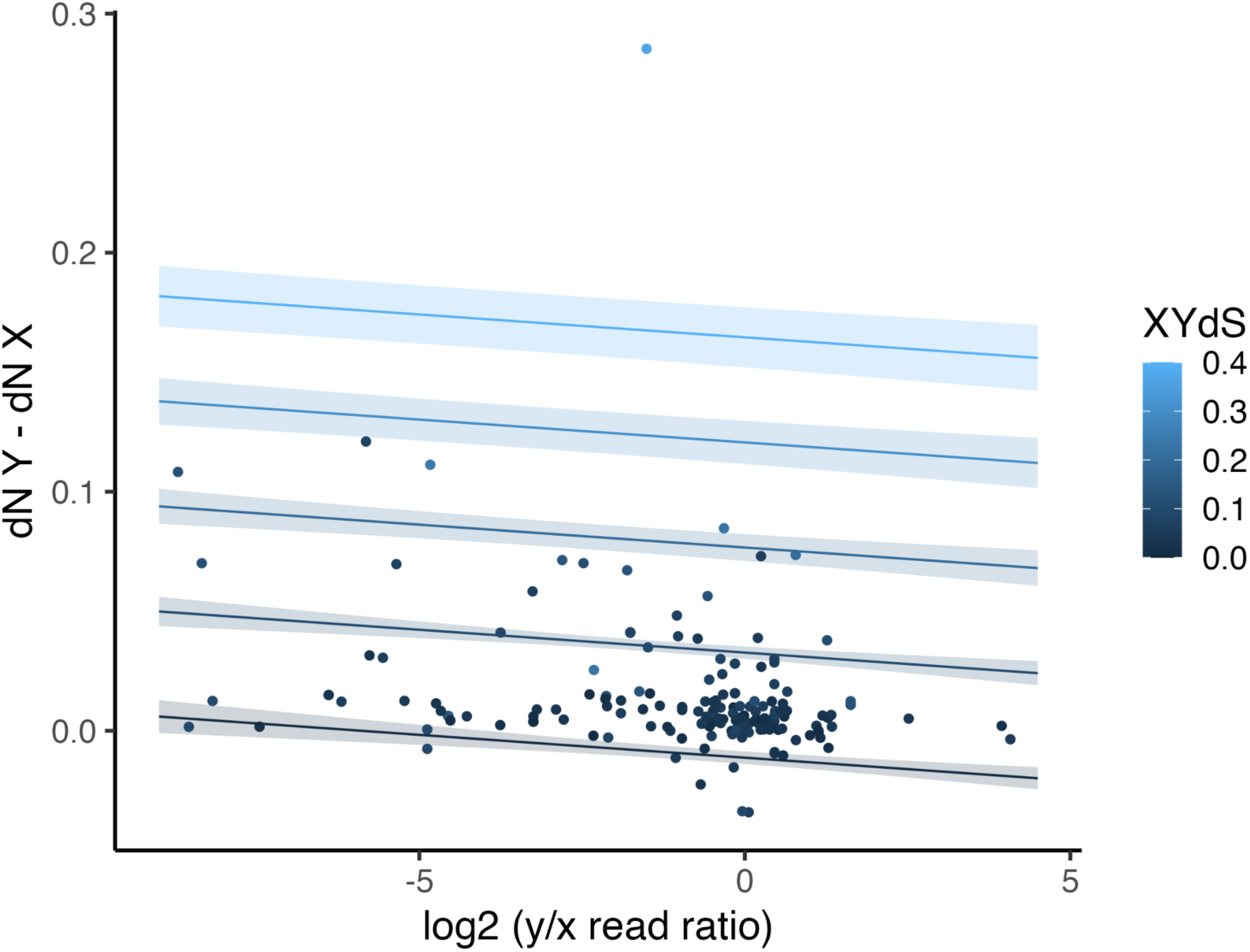
Linear regression of Y/X read ratio and *d_N_* difference between X and Y gametologs. Model: ∼ log2(y/x read ratio) + *d_N_* Y-*d_N_* X + d_S_, *p*-value: < 2.2e-16, Adj *R*^2^ = 0.54. Light to dark blue lines represent different levels of baseline synonymous substitution for gametologs.

### Unique and conserved hemizygous genes identified between Rumex hastatulus cytotypes

Out of 1031 orthologs shared between the sex-linked region (excluding the PAR) of the X chromosome of *R. hastatulus* XY subtype and hermaphroditic relative *R. salicifolius,* we identified 360 putatively lost genes which did not have an annotated ortholog on the *R. hastatulus* Y chromosome, 347 of which were confirmed to be lost via BLAST (Table 2). This ∼33.6% percent of gene loss is very similar to the degree of Y gene loss (∼30%) observed in the old sex-linked region of the XYY subtype from our previous study (Sacchi & Humphries et al. 2024). Of those 347 genes lost on the Y, 64% were defined as completely lost as there were no blast hits to the genomic sequence of the Y chromosome, while 37% can be considered partially lost, as less than half of the sequence was aligned to the Y. One hundred and eighty-three (183) hemizygous genes (approx. 53%) identified in the XYY subtype (Sacchi & Humphries et al. 2024), were also confirmed to be hemizygous from this analysis in the XY subtype, with enrichment of those that are completely lost (Table 2). Three hundred and fifty-two (352) Y genes remain on both cytotypes, excluding the PAR, which is rich in shared genes, as expected. The proportions of present, completely missing and partially missing genes in the XY and XYY subtypes were not uniform (*X*-squared = 466.98, df = 4, *p*-value < 2.2e-16) (Table 3). Notably, 92 genes were completely absent from the Y-linked region of the XY subtype and present in the XYY subtype, and 34 genes were completely absent from the XYY older Y-linked region while present in the XY subtype. This finding suggests an unequal pattern of uniquely lost Y genes between each cytotype, with more unique gene loss taking place in the XY subtype.

**Table 2.**
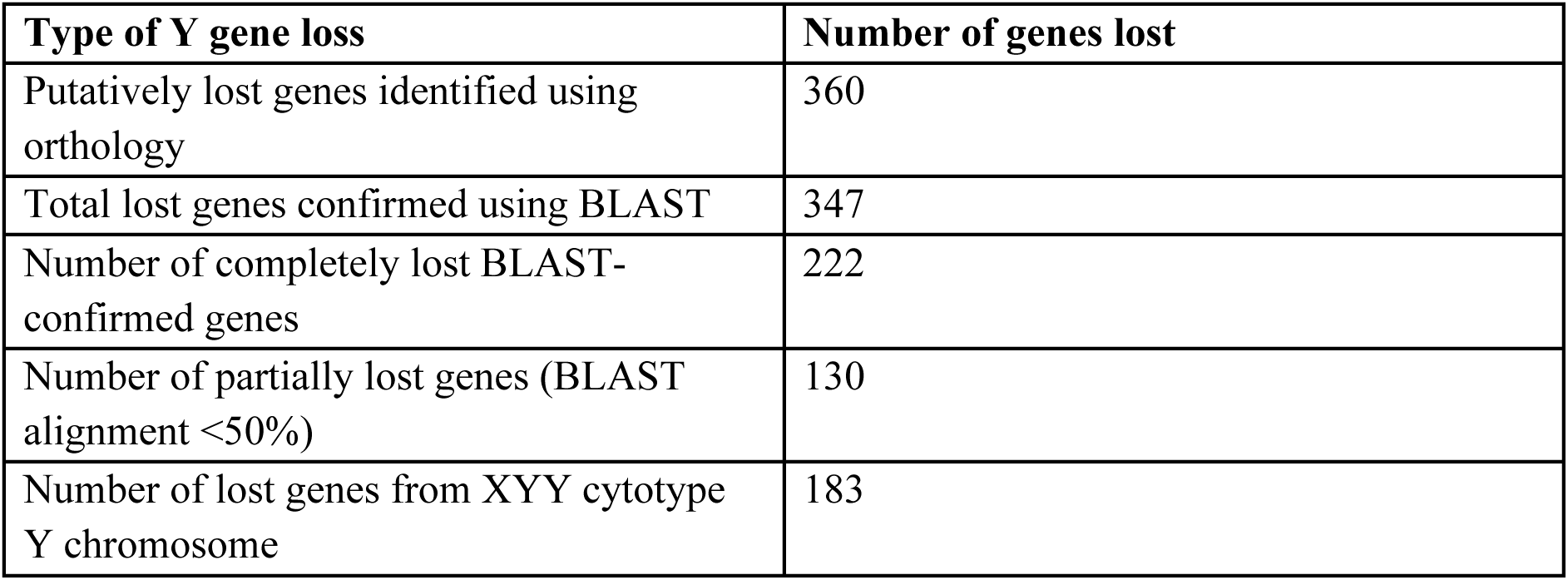
Number and types of hemizygous genes lost from the Y sex-linked region of *Rumex hastatulus*.

**Table 3.**
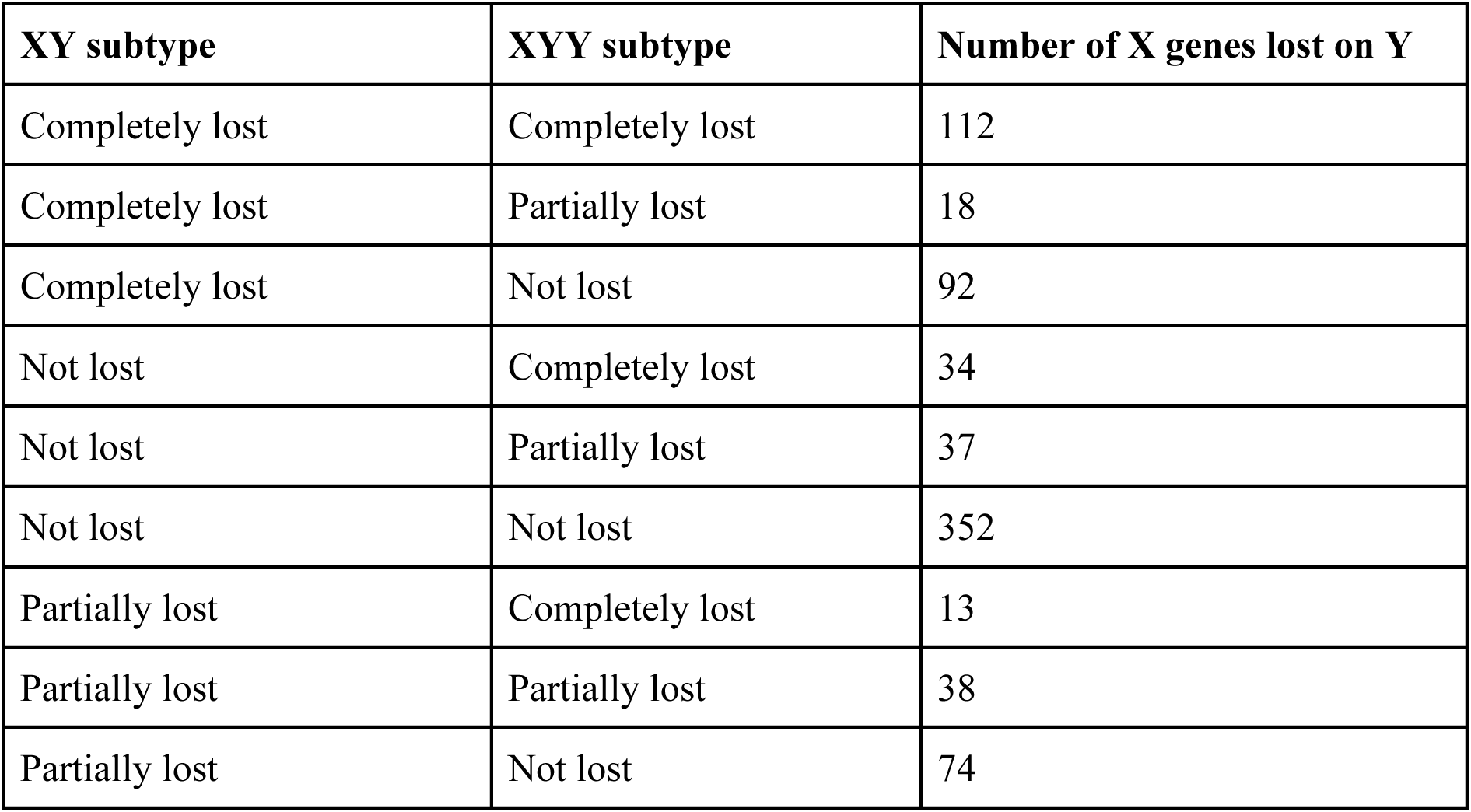
Comparison of completely and partially lost genes with orthology between XY and XYY subtypes of *Rumex hastatulus*.

| XY subtype | XYY subtype | Number of X genes lost on Y |
| --- | --- | --- |
| Completely lost | Completely lost | 112 |
| Completely lost | Partially lost | 18 |
| Completely lost | Not lost | 92 |
| Not lost | Completely lost | 34 |
| Not lost | Partially lost | 37 |
| Not lost | Not lost | 352 |
| Partially lost | Completely lost | 13 |
| Partially lost | Partially lost | 38 |
| Partially lost | Not lost | 74 |

### No evidence of widespread dosage compensation

If partial or complete dosage compensation had occurred in *R. hastaulus*, male expression of hemizygous or silenced X-linked genes (defined as those with significantly reduced Y expression) should significantly exceed 50% expression of females. In the absence of dosage compensation, X-linked gene expression in males is expected to be halved. A comparison of mean X gene expression between male (*n*=75) and female (*n*=74) samples for both hemizygous genes and Y silenced genes revealed no widespread patterns suggesting dosage compensation. In general, nearly all of both types of genes fell very close to the 50% expression expectation on the X chromosome (Figure 5). There were a handful of genes with Y overexpression relative to X expression and they displayed no evidence of dosage compensation when comparing male and female X expression (Supplementary Figure 3). Statistical tests for differential expression of X-linked genes between males and females also suggested no widespread dosage compensation. We used an FDR-corrected *p*-value cutoff of 0.1 and an absolute log2 fold change cutoff of 1. Eighteen percent (113/612) of hemizygous and shared X-Y gametologs were significantly upregulated in females and only one gene was upregulated in males. To further explore evidence for dosage compensation, we doubled read counts of X-linked genes in males only, to test whether two-fold X expression in males is significantly higher than female X expression. Using this approach, five (<1%) X genes were significantly upregulated in males, which indicated possible dosage compensation, whereas the rest were not significantly different from females (Supplementary Table 3). Thus, the overall patterns of X expression in males and females of *R. hastaulus* combined with statistical tests reveal minimal evidence of dosage compensation in this species.

**Figure 5.**
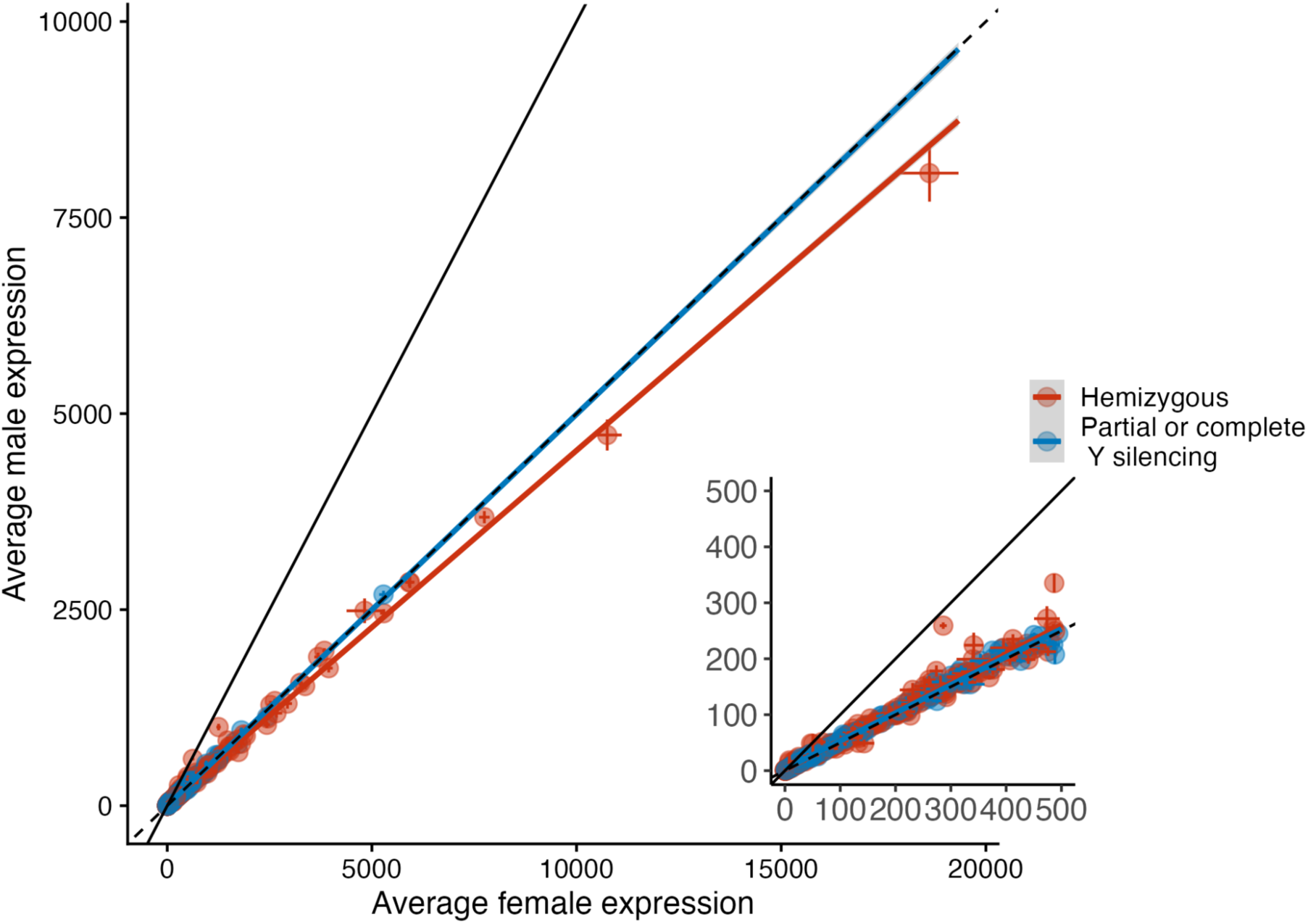
Comparison of X chromosome gene expression between males versus females of *Rumex hastatulus*. Each point represents a single X chromosome gene present in both males and females. Gene expression was averaged across 75 male samples and 74 female samples. Hemizygous genes (red), have no Y gametologs present in the male genome and are indicative of gene loss on the Y. Partially or completely silenced genes (in blue) have significantly decreased Y/X expression ratios (log2FC <1, FDR adjusted *p-*value <0.1). Error bars represent the standard error of mean gene expression for each sex. The solid black line has a slope of 1 and represents equal male and female X expression ratios, which is expected under widespread dosage compensation. The dashed line has a slope of 0.5, which is expected when X expression in males is half that of females, as expected in the absence of widespread dosage compensation.

## Discussion

We used a large, highly replicated transcriptomic dataset and a genome assembly that includes fully-phased X and Y chromosomes to examine the regulatory changes that have occurred in the first 10 million years of sex chromosome evolution in the XY subtype of *Rumex hastatulus*. We found that gametolog-specific expression is biased toward increased X expression relative to Y expression (Figure 1). Our results are consistent with widespread loss of expression in Y genes during the early stages of Y chromosome degeneration. Under the regulatory hypothesis of Y degeneration (Lenormand and Roze 2022), an early and asymmetric increase in X-linked cis-regulatory strength relative to that of Y cis-regulators is predicted. Additionally, this model predicts the accumulation of deleterious mutations on the under-expressed Y genes, leading to further Y expression loss and mutation accumulation. We uncovered a negative relationship between Y/X expression ratio and the nonsynonymous substitution rate on the Y chromosome (Figure 4), which is consistent with the regulatory degeneration model. However, we did not observe early dosage compensation evolution, as the model predicts in hemizygous and silenced genes (Figure 5). While some conditions of the regulatory divergence model are met, a critical component, early dosage compensation, is absent in this system. The lack of early dosage compensation suggests that regulatory divergence may be the result of other forces. For example, selective interference resulting in TE accumulation disrupting Y-linked gene regulation, or adaptive silencing following mutation accumulation (Bachtrog 2013; Crowson et al. 2017).

Regulatory models of sex chromosome degeneration predict early evolution of gametolog-specific expression, where Y expression is lower than X expression, due to differences in cis-regulatory strength. We found a pattern of decreased Y/X expression (Figure 1) in a sex chromosome pair that diverged less than 10 MYA. The observed expression changes taking place in the early phases of sex chromosome evolution are consistent with some aspects of recent regulatory models of degeneration, which predict expression divergence to occur before and alongside widespread deleterious mutation accumulation (Lenormand and Roze 2022). These patterns of decreased Y/X expression are consistent with previous transcriptome-only-based approaches in *R. hastatulus* and other members of the genus (Hough et al. 2014; Crowson et al. 2017). Our methods reduce the likelihood of mapping bias causing false signals of gametolog-specific expression, and provide a more confident estimate of gametolog-specific expression at the gene level compared to transcriptome-based studies. Additionally, our highly replicated data set improves our ability to precisely measure and identify significant gametolog-specific expression patterns. Since our filtering process excluded genes with genomic mapping bias and genes with high Y mis-mapping, this led to a significant reduction in gametologs we could confidently analyze, which may affect our results. A less conservative filter based on the percentage of female reads mapping to the Y, instead of the total number of reads, may increase the number of gametologs in the final data set.

In *Silene latifolia*, recent work yields similar patterns, finding the median of Y/X gametolog expression below 1, and correlated with increased genic methylation on the Y (Moraga et al. 2025). Currently, the mechanism for Y/X underexpression in *R. hastatulus* remains unknown, and future work is necessary to identify possible cis-regulatory variation or epigenetic factors associated with increased X and/or decreased Y expression. Whether any related cis-regulated variants are under selection will be essential for untangling why we observe decreased Y/X expression, i.e. to disentangle the role of drift-induced weakening of cis-regulatory sequence on the Y with upregulation in the X. Nearby transposable elements and methylation patterns have not yet been identified in the XY subtype; however, in the XYY subtype there is evidence for considerably more TEs in and near genes in the older Y chromosome (homologous to the Y in this present study) than anywhere else in the genome (Sacchi & Humphries et al. 2024).

We also observed a small number of gametologs with significantly increased Y expression relative to X expression (Figure 1). In other sex chromosome systems, there are several examples of increased Y expression or amplicon number, especially for genes with specialized male function (e.g. spermatogenesis) (Shaw and White 2022). While only 1:1 orthologs were examined, future work will identify orthologs with copy number variation between the sex chromosomes and determine whether highly expressed Y genes or copy number variants (CNVs) have male-specific functions, as observed in other young sex chromosome systems (Bachtrog et al. 2019).

We observed a significant loss of Y chromosome genes when compared to homologous genes present on the X chromosome and an outgroup. This observation is similar to the rate of gene loss observed in the XYY subtype of *R. hastatulus* (Sacchi & Humphries et al. 2024), which shares the same old sex-linked region despite having diverged approximately 150,000 years ago. The proportion of Y gene loss is similar in both *R. hastatulus* subtypes (∼30%) and is less extensive than Y degeneration observed in other plant sex chromosomes, such as *Silene latifolia* (Moraga et al. 2025), where estimated gene loss is 58%. Over the course of ∼150,000 years, there are some differences in the type of gene loss observed on the Y chromosome. While many genes are completely lost in both subtypes, suggesting an older evolutionary origin prior to divergence, hundreds are uniquely lost only in one subtype (Table 3). Specifically, there are nearly three times more unique Y gene loss events in the XY subtype than the XYY subtype and two times more partial gene loss events unique to the XY subtype. Given that the non-recombining region is larger in the XYY subtype (296 Mb X and 503 Mb Y1+Y2, versus 220Mb X and 414Mb Y), we might expect increased mutation and gene loss due to the Hill-Robertson effect (Hill and Robertson 1966); however, we observed the opposite: greater Y gene loss in the smaller sex-linked region. This result may be due to stochastic differences in gene loss events occurring post-divergence during the last 150,000 years. These differences could either be due to further rapid degeneration of an already-degenerating sex chromosome, or population-level variation in gene loss, only a small part of which is reflected in the reference individual. A pangenome approach that compares gene presence-absence variation across individuals in the sampled *R. hastatulus* population may help fully resolve these patterns.

Variation in median synonymous substitution rate (*d_S_*) across the sex-linked region suggests two or possibly three evolutionary strata, the youngest with a median *d_S_* well below 0.03, and the oldest with median *d_S_* approaching 0.04 (Figure 3b). The high *d_S_* region immediately proximal to the pseudoautosomal region was followed by a sharp drop in *dS*, which may be due to a rearrangement. The presence of a possible inversion between the sex-linked region and the PAR is also supported by Hi-C contact map patterns (Supplementary Figure 5). Genes with significant gametolog-specific expression were found in regions with higher *d_S_*, which suggests that differences in gametolog-level expression accumulate over time following recombination suppression. While here we use *d_S_* as an estimate of time since divergence, low *d_S_* can also result from processes such as gene conversion, which has been suggested to protect essential Y genes from degeneration (Rosser et al. 2009; Betrán et al. 2012).

In our study Y expression loss is significantly associated with a greater non-synonymous substitution rate on the Y gametolog. While these patterns are consistent with the degeneration by regulatory evolution hypothesis (Lenormand et al. 2020), mutation accumulation caused by selective interference or adaptive silencing may also result in decreased Y expression correlated with nonsynonymous mutations (Crowson et al. 2017; Shaw and White 2022). This is because it is difficult to discern from these data whether this accumulation of non-synonymous mutations is an effect of decreased Y expression, a cause, or both processes. Previous reference transcriptome-based work in *R. hastatulus* found that genes on the X under positive selection were not more likely to be silenced on the Y (Beaudry et al. 2017), suggesting that important genes may not be silenced in response to mutation accumulation. In the related *R. rothschildianus* there is no correlation between X-Y *dn*/*ds* and Y/X expression, in contrast with prior predictions of active Y suppression, nor is there a correlation between positive selection on X genes and the degree of Y silencing (Crowson et al. 2017). Thus overall, our results are consistent with the prediction of greater deleterious mutation accumulation on Y-linked genes experiencing expression loss, but further population genomic analysis would be important to test this further.

We did not observe evidence of widespread dosage compensation in hemizygous and Y-silenced genes (Figure 5). While there are a small number of X genes in males that may be significantly upregulated to female levels, female X expression is approximately twice that of male X expression for most hemizygous and silenced genes. Several selective interference models of chromosome degeneration predict that dosage compensation arises as the Y becomes sufficiently degenerated, leading to selection for increased X expression and silencing of the Y chromosome (Charlesworth 1978; Orr and Kim 1998). On the other hand, dosage compensation is predicted to arise early on if Y degeneration and the spread of recombination suppression is driven by regulatory evolution (Lenormand et al. 2020; Lenormand and Roze 2022). Early dosage compensation is a critical component of the regulatory divergence model, since dosage compensation facilitates fixation of the initial recombination-suppressing inversions (Lenormand and Roze 2022).

Our results suggest that degeneration, both in terms of Y under-expression and gene loss, occurred during the 5-10MYA of recombination cessation; however, widespread dosage compensation is not apparent. In some plant species that have been investigated what is termed ‘partial’ dosage compensation (e.g. *Silene latifolia*) dosage compensation-like expression patterns were observed in approximately half of the hemizygous genes *-* potentially mediated by an enrichment of epigenetic silencing in one female X chromosome (Muyle et al. 2022; Papadopulos et al. 2015). In our study only a handful of genes that we tested appeared to have male:female X expression ratios that are suggestive of dosage compensation and a widespread pattern was not observed. A lack of complete dosage compensation is not unique to plants or animals. Widespread transcriptional dosage compensation is not observed in birds or platypuses, however recent work suggests that extensive post-transcriptional regulation mediates dosage compensation at the proteomic level in these taxa (Lister et al. 2024). It is possible that differences in gene dosage at the genomic and transcriptional levels in plants, such as *R. hastatulus*, are compensated via other mechanisms. Additionally, it is possible that hemizygous genes are under less selective constraint from the outset, or are less dosage sensitive, as may be the case in *R. rothschildianus* where lower expressed or less constrained genes are more likely to be hemizygous (Crowson et al. 2017). In either case, it appears unlikely that early dosage compensation was an important contributor to the fixation of recombination-suppressing inversions or expansion of the sex-linked region of *R. hastatulus*. The extremely large non-recombining sex-linked region may be due to the presence of pre-existing low recombination, particularly in male recombination (Rifkin et al. 2022).

Our study identified widespread Y expression loss during the first 10 million years of sex chromosome loss in the dioecious plant *Rumex hastatulus.* This finding aligns with predictions of the regulatory model of sex chromosome degeneration, but the patterns of increased non-synonymous substitution rate in Y genes with lower gametolog-specific expression could also be a result of selective interference-based processes. Dosage compensation was not observed at this timescale, despite the complete loss and silencing of hundreds of genes, which is more consistent with classical models of sex chromosome degeneration where dosage compensation evolves later in the process of degeneration. The lack of early dosage compensation suggests that regulatory divergence may not be playing a role in recombination suppression and degeneration, despite the overall pattens of X-Y expression divergence. Our results add to a growing body of work investigating expression and degeneration patterns in young sex chromosomes and these likely hold the key to understanding the order of events and types of processes resulting in Y chromosome degeneration.

## Data Availability

Assembled haplotype-resolved reference genome (sequence and annotations) for the *Rumex hastatulus* XY cytotype male are available at the Borealis data repository https://doi.org/10.5683/SP3/FRHNO4 and NCBI under Bioprojects PRJNA1367036 and PRJNA1367037. Analysis scripts are available at https://github.com/bmsacchi/rumex_XY_expression.

