## Supplemental Materials for "Widespread loss of Y expression in the absence of transcriptional dosage compensation in *Rumex hastatulus*"

**Supplementary Table 1. Assembly statistics for each haplotype of *Rumex hastatulus***

| Assembly | Total length | Scaffold N50 | Scaffold L50 | N90 | L90 | BUSCO Score (%) | Contig N50 | Contig L50 |
|---|---|---|---|---|---|---|---|---|
| Haplotype A | 149742293 | 315705859 | 2 | 173846016 | 5 | 97.7 | 7,699,624 | 61 |
| Haplotype B | 1863807873 | 465013087 | 2 | 179199150 | 5 | 99.2 | 5,565,000 | 93 |

**Supplementary Table 2. Number of X-Y gametologs retained after various filtering steps.**

| **Filtering category** | **Number of X-Y gametologs remaining in expression dataset** |
|---|---|
| No filtering | 2324 |
| Removing genes in pseudoautosomal region | 2255 |
| Removing genes with significant Y mismapping in females (RNA data) | 1476 |
| Removing genes with significant Y mismapping in females (genomic data) | 912 |
| Removing genes with significant mapping bias in genomic data | 568 |
| Removing genes with total expression < 20 reads across all samples | 267 |

| **Supplementary Table 3. Differentially expressed X genes in males versus females of *Rumex hastatulus*** | | |
|---|---|---|
| **Expression type** | **Differential expression between males and females (X male vs. XX female)** | **Differential expression between males and females, X read counts doubled (2X male vs. XX female)** |
| Male upregulation | 1 | 5 |
| Female upregulation | 113 | 0 |
| Non-significant: | 498 | 607 |
| Total | 612 | |

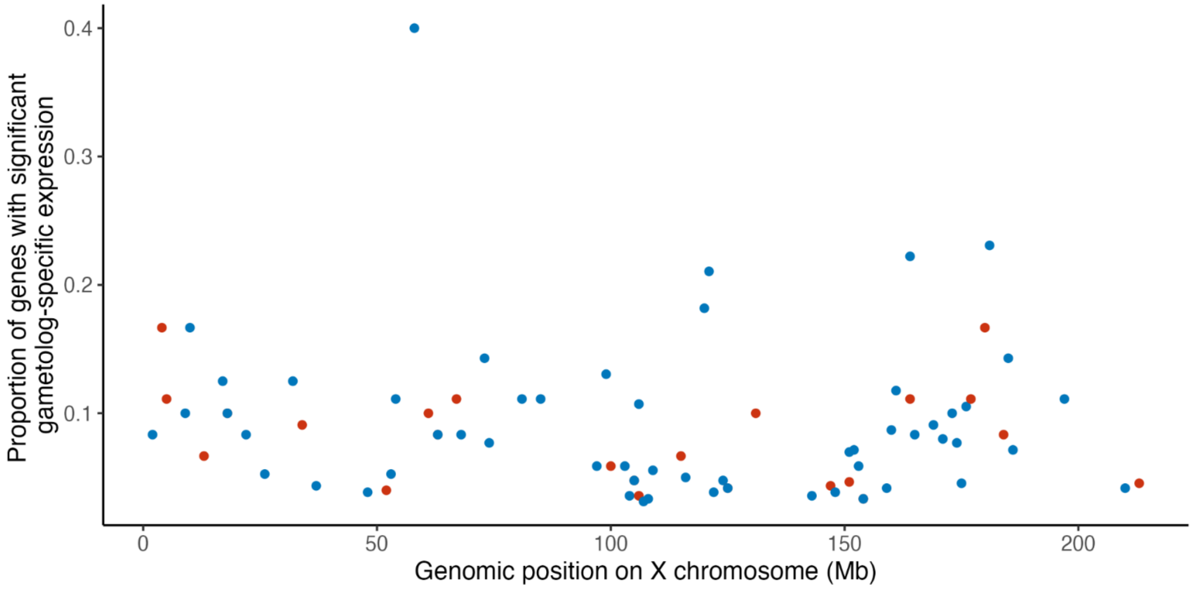


**Supplementary Figure 1.** Proportion of genes in ***Rumex hastatulus*** with significant (p, log2fc >1) gametolog-specific expression in 1mb windows (relative to X chromosome position). Blue points show Y/X expression <1, red points show Y/X >1.

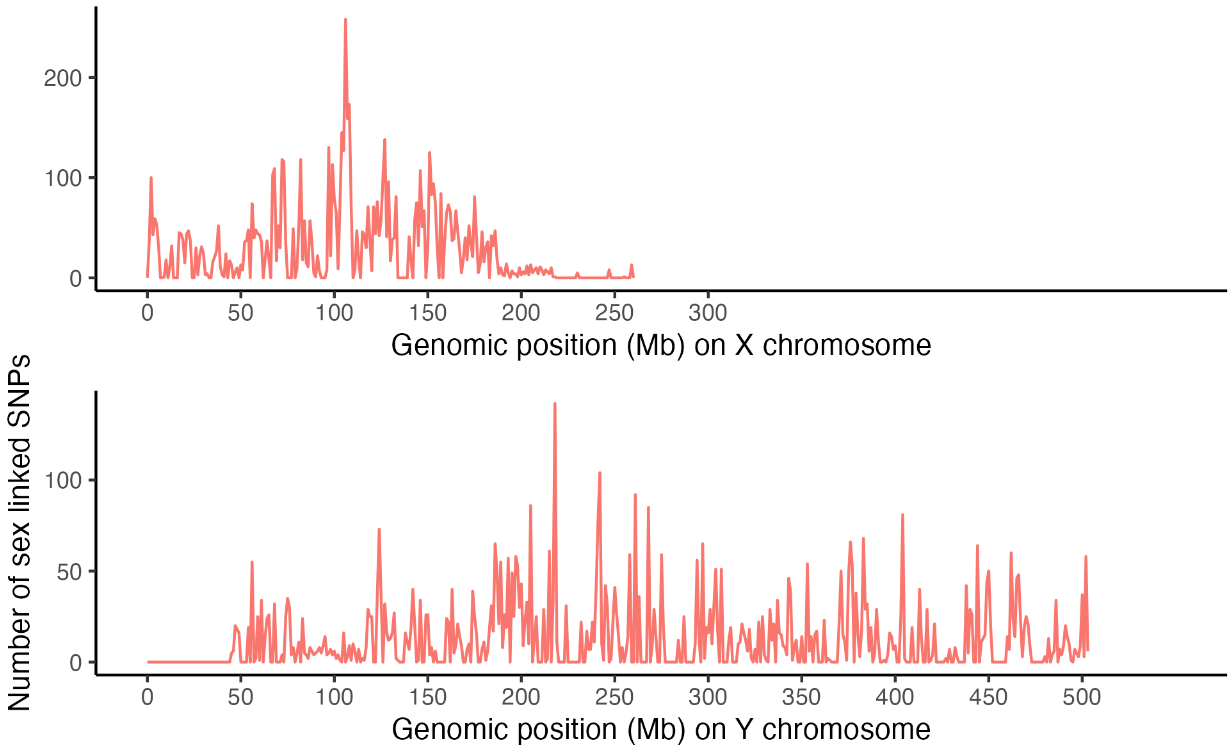


**Supplementary Figure 2.** Density of sex-linked SNPs on the X and Y chromosomes of ***Rumex hastatulus*** per 1MB window.

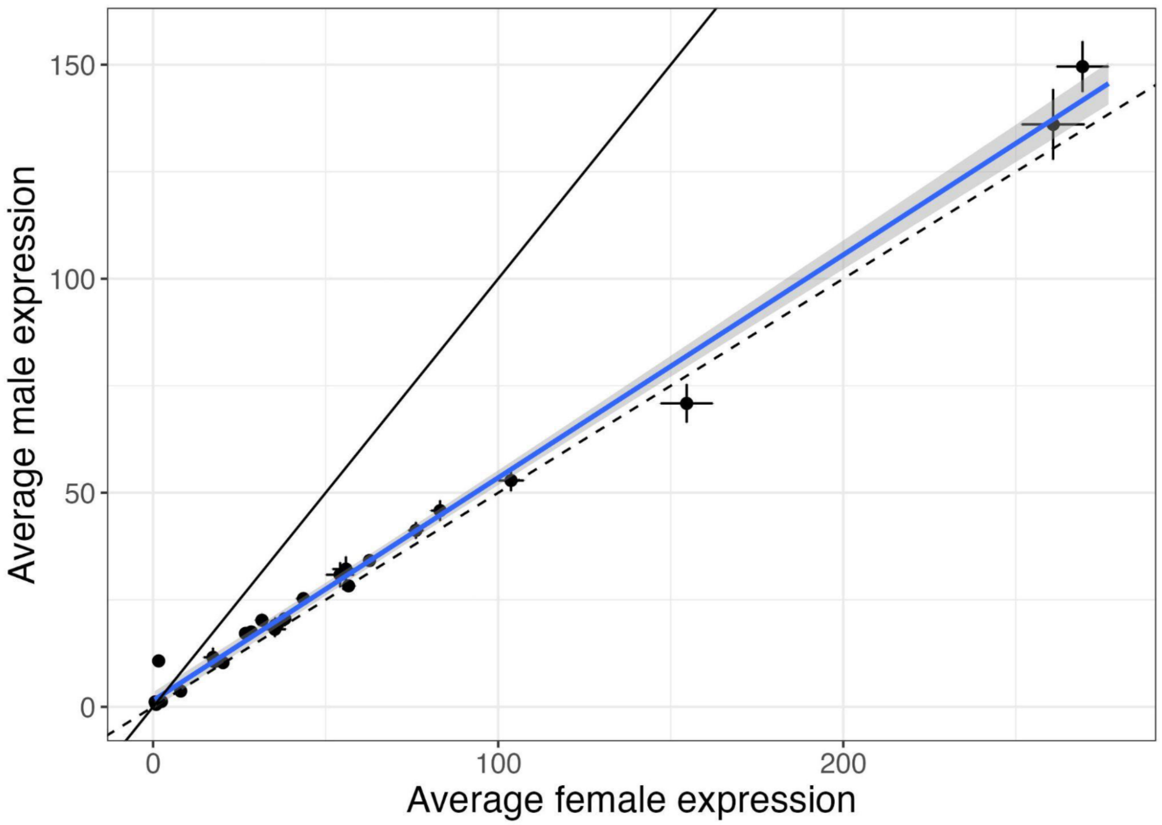


**Supplementary Figure 3**. Average male vs. female expression for genes with Y/X overexpression in ***Rumex hastatulus***. Each point represents a single X chromosome gene present in both males and females. Gene expression is averaged across 75 samples in males and 74 in females. Points represent genes where Y expression is greater than X expression, indicating Y overexpression (log2FC > 1, FDR adjusted *p*-value <0.1). Error bars represent the standard error of mean gene expression for each sex. The solid black line has a slope of 1 and represents equal male and female X expression ratios, which is expected under widespread dosage compensation. The dashed line has a slope of 0.5, which is expected when X expression in males is half that of females, as expected in the absence of widespread dosage compensation.

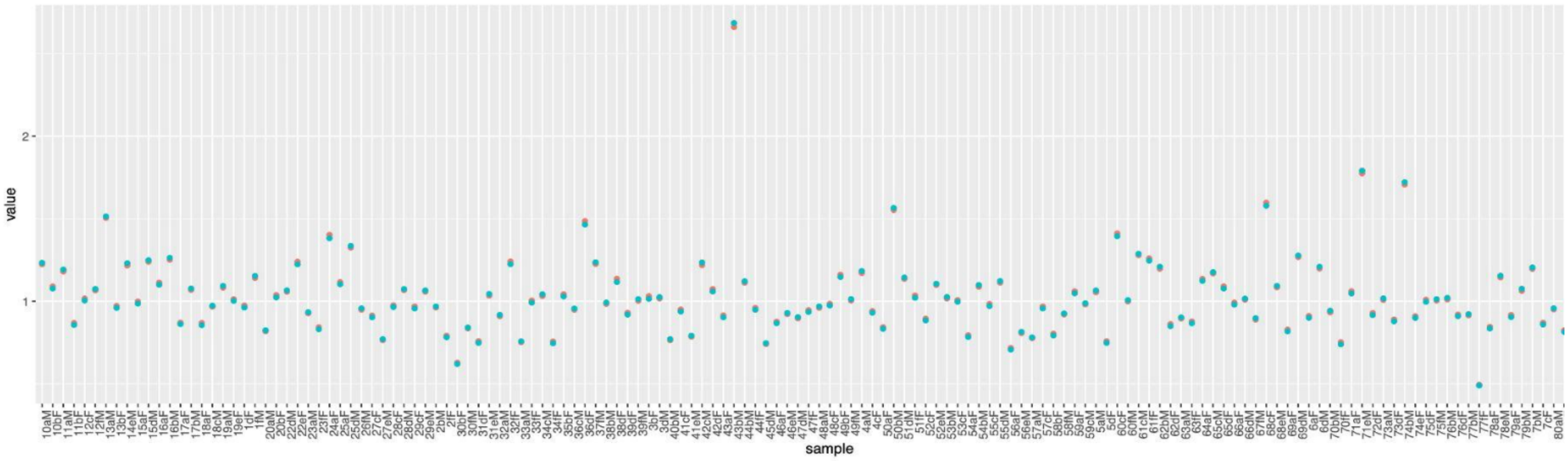


**Supplementary Figure 4**. Comparison of DESeq library size normalization factors for each sample of ***Rumex hastatulus*** when computed with sex chromosomes and autosomes (red) and when computed with autosomes only (turquoise).

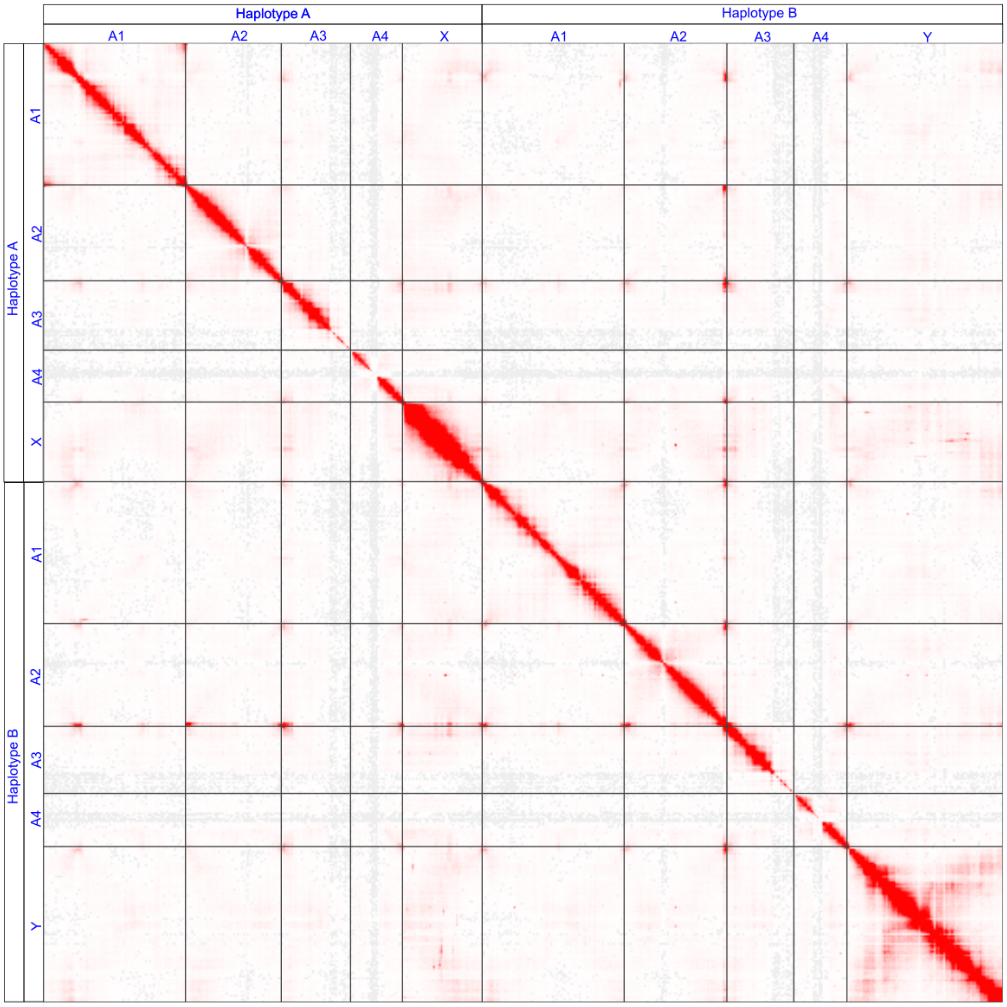


**Figure 5**. Omni-C contact maps for *Rumex hastatulus* XY cytotype. The featured assembly contains all chromosomes from each haplotype.

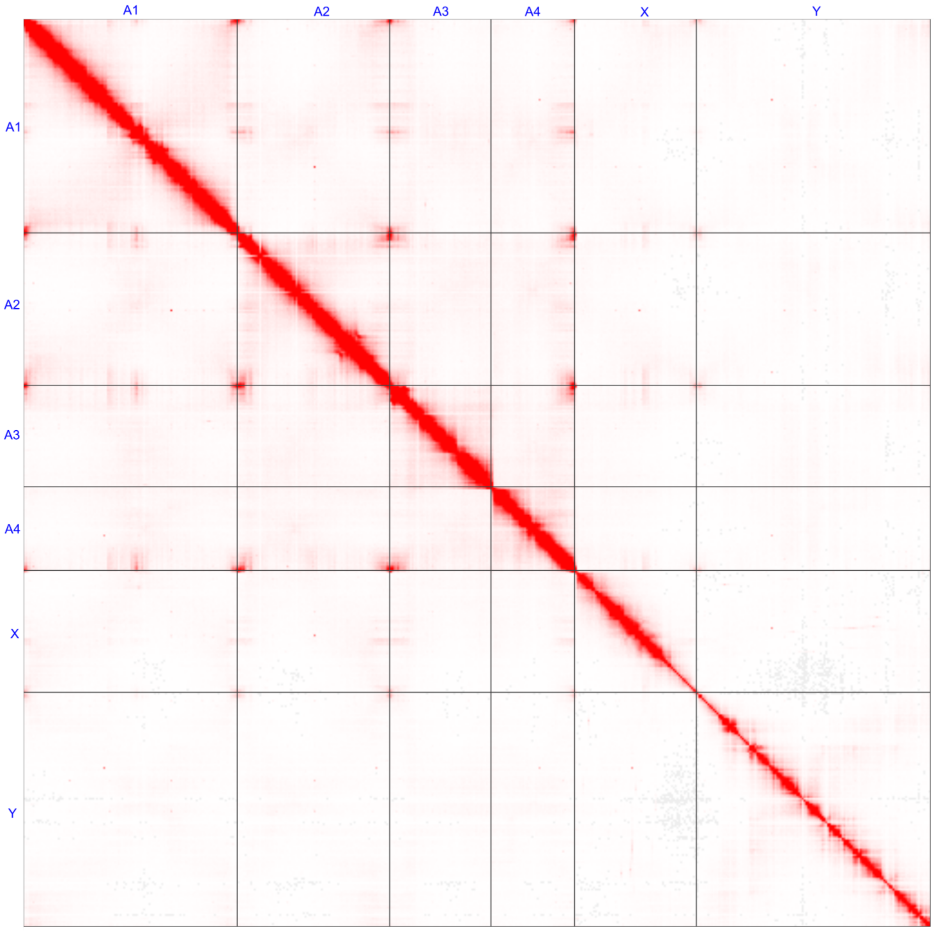


**Figure 6.** Omni-C contact maps for *Rumex hastatulus* XY cytotype. The featured assembly contains the Y chromosome and PAR, all autosomes from the Y-bearing haplotype (haplotype B), and the X chromosome (PAR removed).

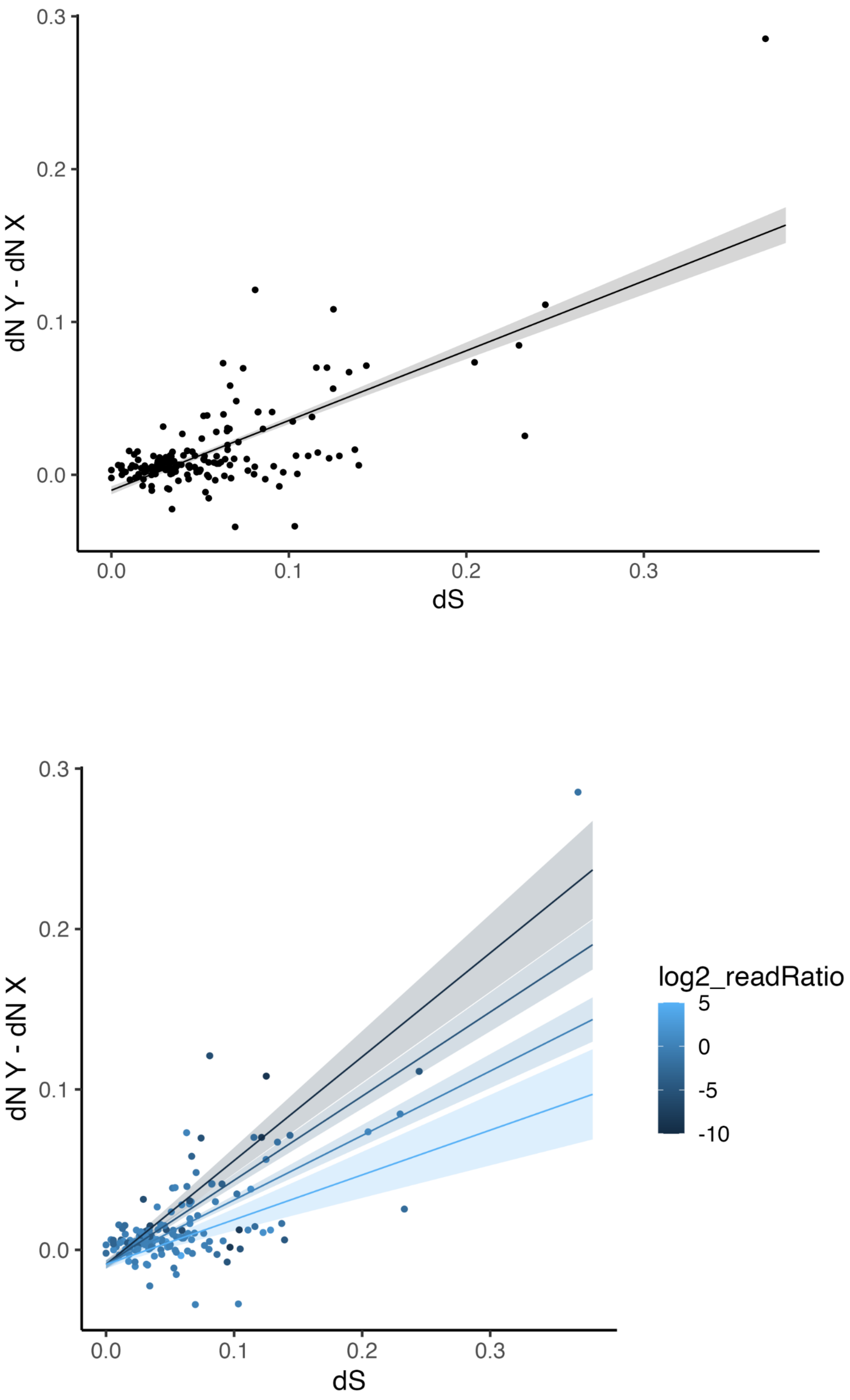


**Supplementary Figure 7**. Relationship between *dN* Y - *dN* X and *dS*. Bottom panel shows the interaction term between log2_readRatio and *dS*.
